# Lossless Immunocytochemistry using Photo-polymerized Hydrogel Thin-films

**DOI:** 10.1101/2020.02.24.963793

**Authors:** Jeong Hyun Lee, Aline T. Santoso, Emily S. Park, Kerryn Matthews, Simon P. Duffy, Hongshen Ma

## Abstract

Immunocytochemistry (ICC), or immunofluorescence microscopy, is an essential biological technique for phenotyping cells in both research and diagnostic applications. Standard ICC methods often do not work well when the cell sample contains a small number of cells (<10,000) because of the significant cell loss that occurs during washing, staining, and centrifugation steps. Cell loss is particularly relevant when working with rare cells, such as circulating tumor cells, where such losses could significantly bias experimental outcomes. In order to eliminate cell loss in ICC protocols, we present a method to encapsulate the cell sample in a photo-polymerized hydrogel thin-film. The hydrogel thin-film is permeable to antibodies and other ICC reagents, thereby allowing the use of standard ICC protocols without modification. The cell sample is physically constrained by the hydrogel at the bottom surface of a standard (unmodified) imaging microtiter plate, thereby enabling the acquisition of high-quality micrographs regardless of the properties of the cell sample or staining reagents. Furthermore, while standard ICC requires several centrifugation steps during staining and washing, our hydrogel encapsulation method requires only a single centrifugation step. This property greatly reduces the time required to perform ICC protocols and is more compatible with robotic platforms. In this study, we show that standard ICC and Cytospin protocols are extremely lossy (>70% loss) when the sample contains less than 10,000 cells, while encapsulating the cells using a permeable hydrogel thin-film results in a lossless ICC process.

## Introduction

Immunocytochemistry (ICC), or immunofluorescence microscopy, is an essential biological assay that uses fluorescence-conjugated antibodies to label cells in order to phenotype them based on protein expression and localization. This assay involves repeated exchange of reagents for cell fixation, permeabilization, blocking, immunostaining, as well as additional buffer washes between each step. When the specimen contains a large number of cells (typically >10^5^ cells per ml), there is sufficient cell density to form a pellet during centrifugation, which enables supernatant removal by pipetting or decanting the fluid. However, when there are fewer cells, the cell density is too low to pellet and many cells may be lost during each supernatant removal step. This issue is particularly important when working with precious samples, where the specimen is limited, or where target cells are rare. For example, detecting circulating tumor cells (CTCs) in the blood of cancer patients^1–4^ or fetal cells in maternal blood^5^, require immunostaining of exceedingly rare cells, where the loss of potential target cells cannot be tolerated.

Numerous modifications of the conventional ICC protocol have been developed to prevent cell loss. One approach is to chemically attach cells on a glass slide coated with an adhesive, such as poly-L-lysine, fibronectin, or Cell-tak^6–8^, and then perform the ICC protocol directly on the glass slide. This approach works well for adherent cells grown in culture, but the adhesives are typically ineffective for primary cells or cultured suspension cells. An alternative approach is Cytospin™, which physically adheres cells to a glass slide using centrifugal force^9,10^. While both primary cells and cells grown in culture can be adhered to a glass slide, this process contributes to significant cell loss. Specifically, when the cell number is relatively small (<10^5^ cells per ml), previous studies have reported losses of >75%^11^. Additionally, Cytospin is a serial process performed one sample at a time, which significantly limits experimental throughput in screening studies^12^. Finally, while Cytospin deposits cells in a confined region on a slide, the deposition area typically requires capturing many microscopy fields to image, which adds to the time required for imaging. Therefore, when the sample contains a small number of cells, concentrating cells in a smaller imaging area can significantly reduce imaging time.

Here, we present a method to prevent cell loss during ICC by encapsulating cells in a hydrogel thin-film. This approach has been used previously by encapsulating cells in low-melt agarose^13^, which forms a hydrogel matrix that is optically transparent and permeable to ICC reagents. However, this approach has not been widely adopted because the viscosity of agarose solutions which prevent the alignment of cells to a precision surface for imaging. Instead, the agarose hydrogel must be sectioned to image the cells from each optical plane. In this study, we present a cell encapsulation material that has lower density than typical cells in order to enable the alignment of cells by centrifugation to a single layer on the bottom surface of a standard and unmodified imaging microtiter plate. This material is permeable to immunoglobulins, optically transparent with minimal coloration and auto-fluorescence, and mechanically robust to withstand repeated washing. Through additional experiments, we show that the lossless ICC process is able to (i) retain and stain 100% of the cell sample, (ii) confine the cell sample into a small area for rapid high-quality imaging, and (iii) can be performed with only a single centrifugation step.

## Results and Discussion

Our general approach is to mix cells in a pre-polymer solution that can be crosslinked into a hydrogel upon ultraviolet(UV) light exposure. To enable lossless ICC, the prepolymer solution must be less dense than cells to allow them to sink to the imaging surface via centrifugation. The hydrogel thin film must have sufficient mechanical strength to withstand normal pipetting. Finally, the hydrogel must be sufficiently thin and porous to allow the diffusion of immunoglobulins in a reasonable timeframe (~1 hour)(Figure 1).

**Fig. 1.**
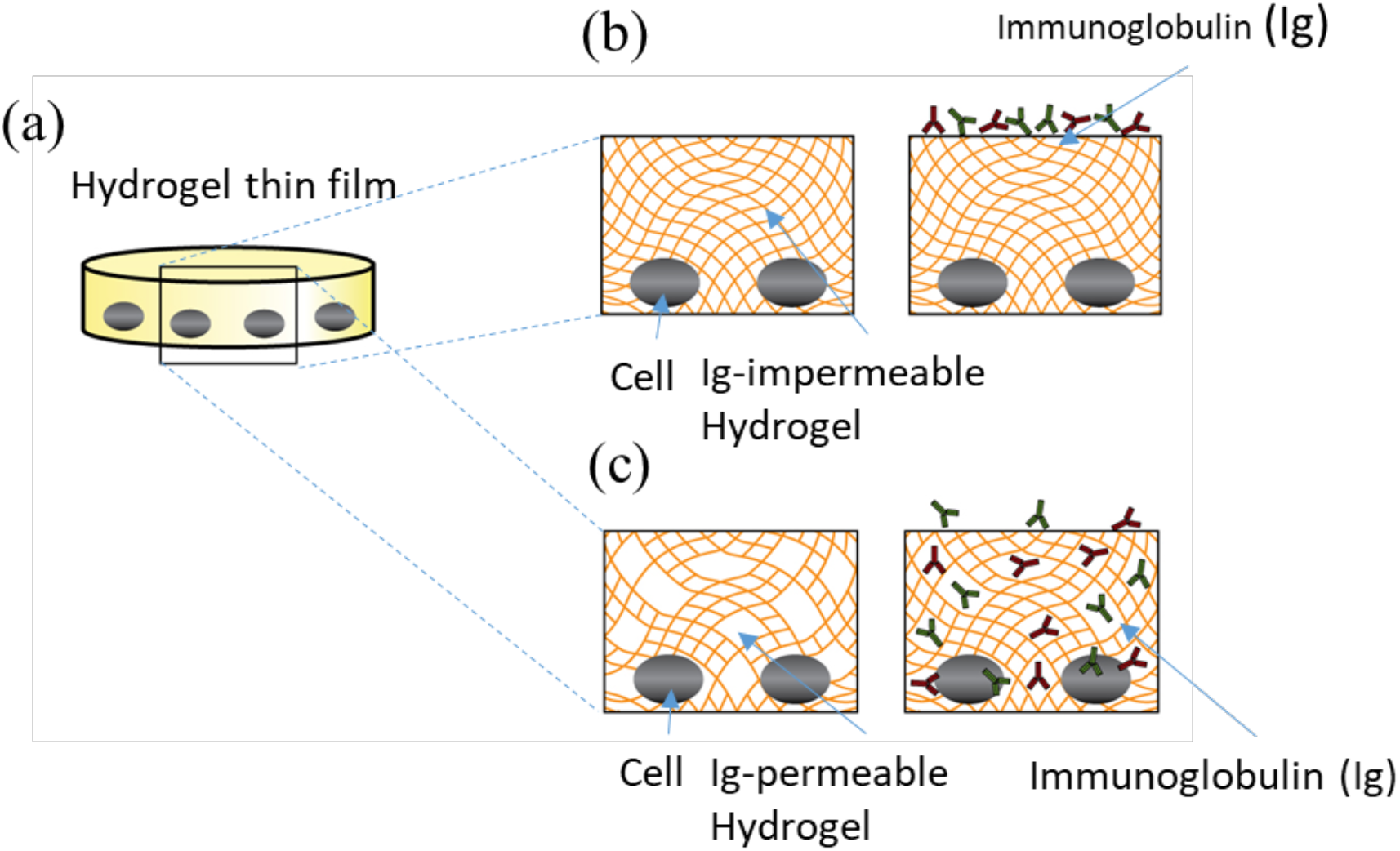
Schematic of the hydrogel. (a) cells encapsulated in photo-polymerized hydrogel. (b) the typical hydrogel is impermeable to immunoglobulins. (c) the hydrogel formulated for immunostaining is permeable to immunoglobulins.

### Density Testing

In order to align cells by centrifugation to a single layer on the bottom glass surface of the imaging microtiter plate, the cell capture solution density must be less than that of typical cells. Given that the lowest density cells are likely to be monocytes, which have a density between 1.067 and 1.077 g/ml^28^, the cell capture solution should be less than 1.067 g/ml in density. Also, the density must be higher than 1 g/ml to sink and encapsulate the cells in bottom plane. We aimed the density of cell capture solution as 1.058 g/ml.

### Mechanical Strength Testing

The mechanical strength of the hydrogel thin-film is important for retaining its structural integrity during pipetting. This property was tested by repeatedly pipetting 40 μl of PBS onto the surface of the photopolymerized hydrogel until signs of structure disintegration, such as cracks, tears, and delamination, began to be observable. The polymerized hydrogel thin-film had sufficient mechanical strength to survive >100 rounds of repeated pipetting.

### Porosity Testing

Immunoglobulins have an estimated size of ~14 nm^14,15^. The porosity of hydrogel thin-film must be sufficiently large to permit diffusion of immunoglobulins to the cell sample in a reasonable amount of time. Conventional method for producing macroporous hydrogels include freeze-drying, solvent casting, and gas forming^16–21^. While these methods have been used in tissue engineering applications to produce hydrogels with >100 μm pores^22–26^, these hydrogels have poor mechanical strength and image quality^27^, which makes them incompatible with immunostaining of embedded cells. We evaluated whether ICC could be performed on cells embedded in lossless hydrogel by embedding 22RV1 cancer cells and then using fluorophore-conjugated antibodies to stain the extracellular EpCAM protein, and the intracellular cytokeratin proteins. The 90% of cells were stained after 1 hour incubation.

### Thickness Testing

In addition to porosity, the thickness of the hydrogel thin-film is important for determining the time required for immunoglobulin diffusion. Immunoglobulin stains are introduced on the top surface of the hydrogel and must diffuse to cells located at the bottom surface of the hydrogel, which interfaces with the glass substrate. The thickness of the hydrogel thin-film can be controlled by the UV light intensity and exposure time. Based on the Beer-Lambert law, UV light intensity diminishes exponentially as it penetrates absorbing material. Therefore, UV light applied at the bottom of the imaging plate polymerizes a hydrogel thin-film with thickness directly controlled by exposure time. We tested hydrogel thin-film formation by exposure using a long-wavelength UV LED (λ=375 nm) for 1, 3, 5, 7, or 10 seconds. We also tested hydrogel thin-film formation by exposure using standard UV gel imaging system (λ=302 nm) for 5, 10, 15, 20, and 30 seconds.

The hydrogel thickness was then estimated by first focusing on a cell along the imaging plate surface and then the top hydrogel surface. The z-position of each focal point was obtained from Nikon NIS-BR software and used to estimate the distance between the two points. Three measurements were performed for each experimental condition. Using an UV LED, exposure for 1 and 3 s failed to form a hydrogel, while 5, 7, and 10 s exposures produced hydrogels with thickness of 100 +/− 20 μm, 500 +/− 20 μm, and 1000 +/− 20 μm, respectively. Therefore, we selected the 5 s as the optimal exposure time for UV LED source because it produced a stable hydrogel while minimizing UV exposure and minimizing hydrogel thickness. When using the UV gel imaging system, exposure <20 seconds failed to form a hydrogel, while 20 seconds exposure produced 100 +/− 20 μm thickness hydrogel and 30 seconds exposure produced 300 +/− 20 μm thickness hydrogel. Therefore, for gel imaging system, we used an exposure time of 20 seconds to minimize hydrogel thickness.

### Immunocytochemistry of Hydrogel Encapsulated Cell Samples

To evaluate this cell fixation method for use in immunocytochemistry, we first mixed the cell sample with the prepolymer solution in a glass-bottom imaging well plate and then centrifuged the well plate to align the cells on the surface of the glass. Next, the hydrogel thin-film is formed using a 5 s UV exposure, at which point, the specimen is immunostained using standard ICC reagents (Figure 2). Since previous efforts to visualize cells in macroporous hydrogels have been limited by image quality^27^, we first evaluated the image quality of hydrogel encapsulated cells by bright field microscopy. While the polymerized hydrogel shows a slight opacity compared to standard buffer, the encapsulated cells were clearly visible under microscopy with no visible degradation in image quality (Figure 3). We then assessed whether the hydrogel impaired immunofluorescence microscopy by staining 22RV1 tumor cells using fluorescence conjugated antibodies specific for EpCAM and pan-cytokeratin (Figure 4). The cells were clearly visible and the absence of background fluorescence indicates that the unbound antibodies were not adsorbed by the hydrogel, but were efficiently washed away. In order to determine whether fluorescence intensity was hindered by hydrogel encapsulation, we measured the fluorescence intensity of cells from the standard ICC sample and the hydrogel encapsulated sample. We measured the mean fluorescent intensity (MFI) from each cell using a 40×40 pixel square area surrounding each cell. The measured MFI did not show a significant difference between the standard ICC sample and hydrogel encapsulated sample (Figure 5). An important consideration was that all staining parameters were optimized using the standard ICC protocol and the conditions did not require re-optimization for hydrogel encapsulated cells. Thus, cell staining could be achieved under standard conditions in <2 hours.

**Fig. 2.**
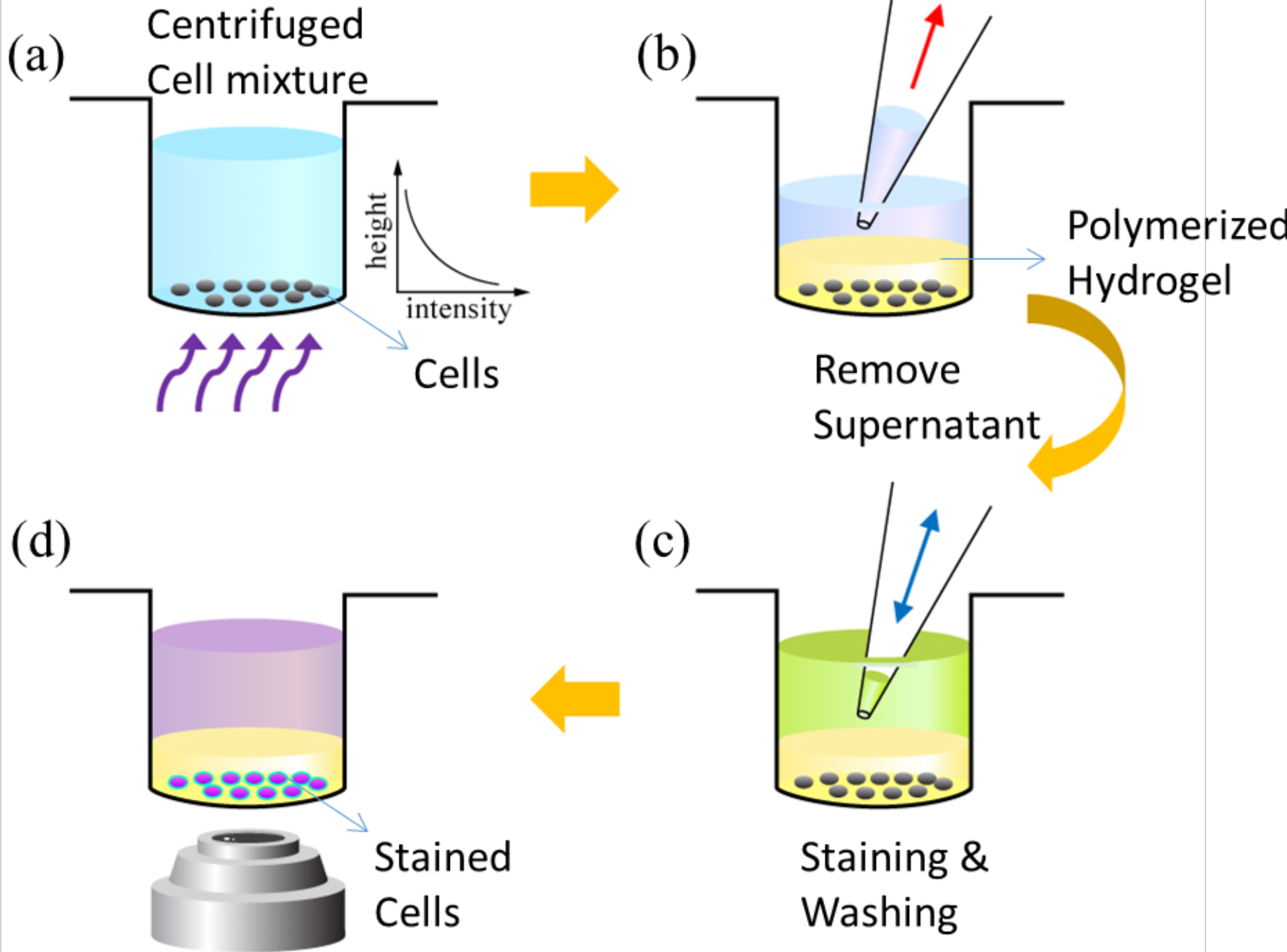
The general approach for lossless immunocytochemistry using hydrogel-encapsulated cells. (a) The cell capture solution and cell suspension is added to a standard (unmodified) glass bottom imaging well-plate. The plate is centrifuged to position the cells at the bottom of the plate and the plate is exposed to UV light for 5 s. (b) The supernatant, along with uncured cell capture solution, is removed from the well by pipetting. (c) Immunostaining steps for fixation, permeabilization, immunostaining, as well as multiple washing steps are performed without additional centrifugation steps. (d) Image acquisition can be performed directly on the imaging plate.

**Fig. 3.**
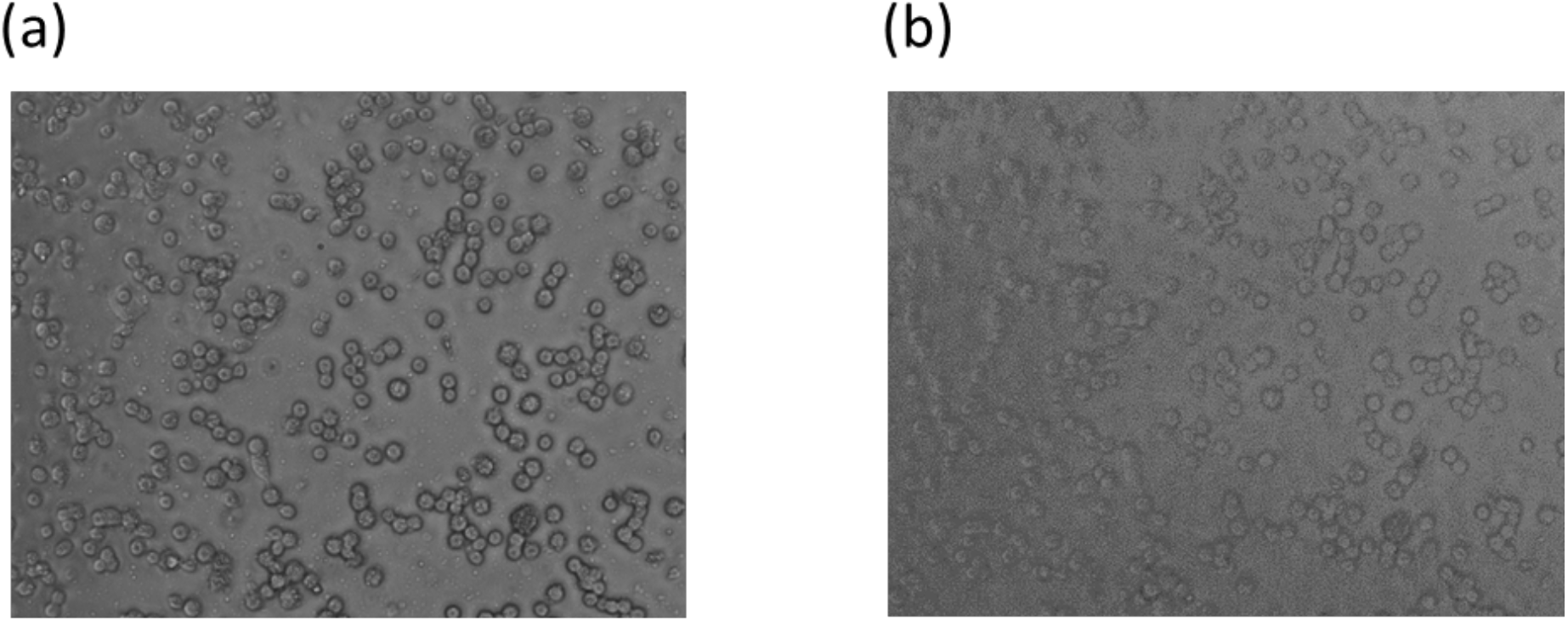
Comparison of bright-field microscopy image of cells encapsulated in hydrogels. (a) before and (b) after photo-polymerization. The polymerized hydrogel shows a slight opacity after photo-polymerization, but there is no significant change in microscopy image quality.

**Fig. 4.**
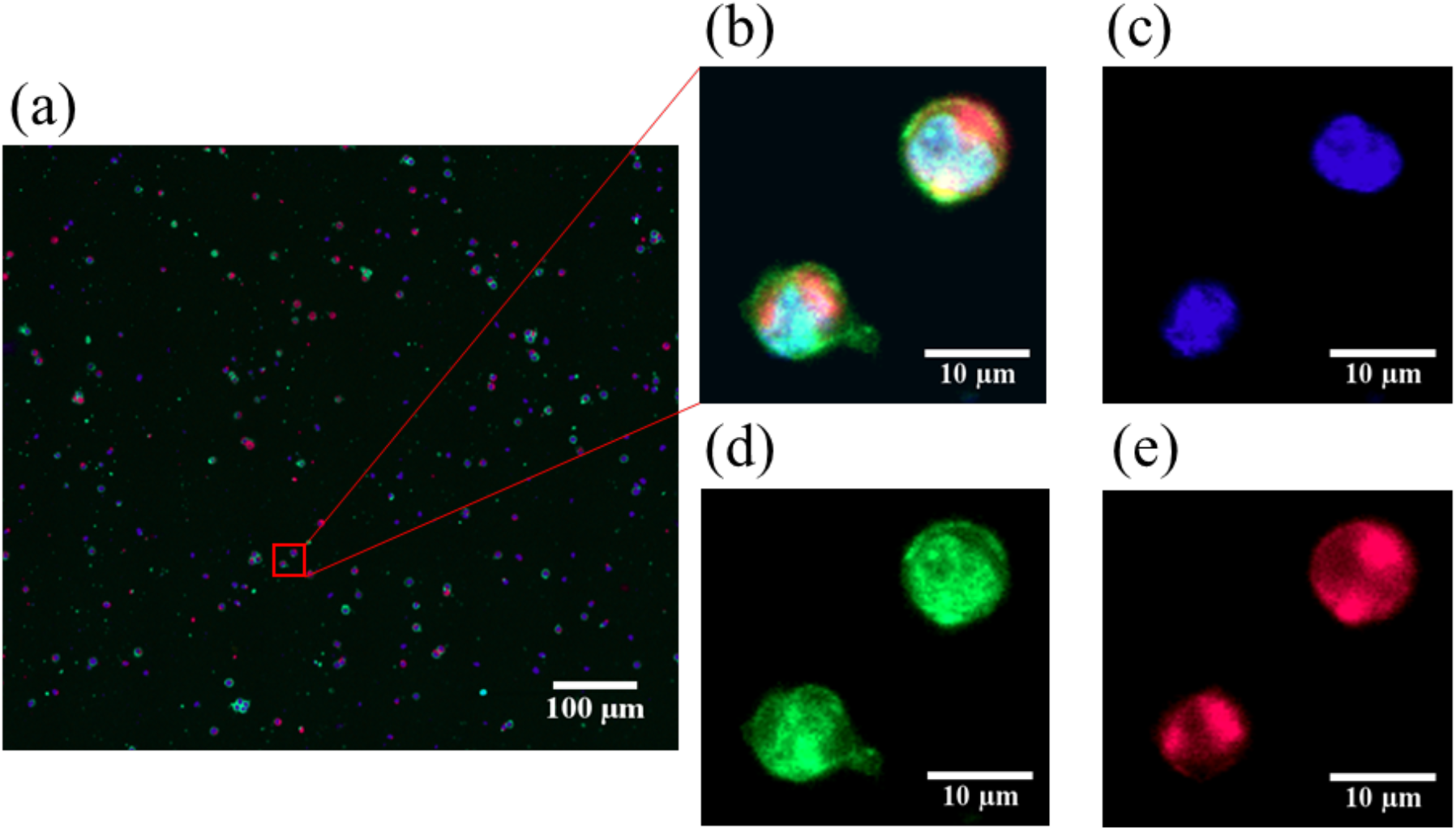
Micrographs of hydrogel encapsulated cells stained with fluorescent antibodies. (a) 40X image from a well in a 384 well microtiter plate where ~1,000 cells are encapsulated in hydrogel and subsequently stained with DAPI (blue), EpCam-Alexafluor-488 (green), Pan-Keratin-Alexafluor-647 (red). (b-e) Close-ups of merged and separated channels.

**Fig. 5.**
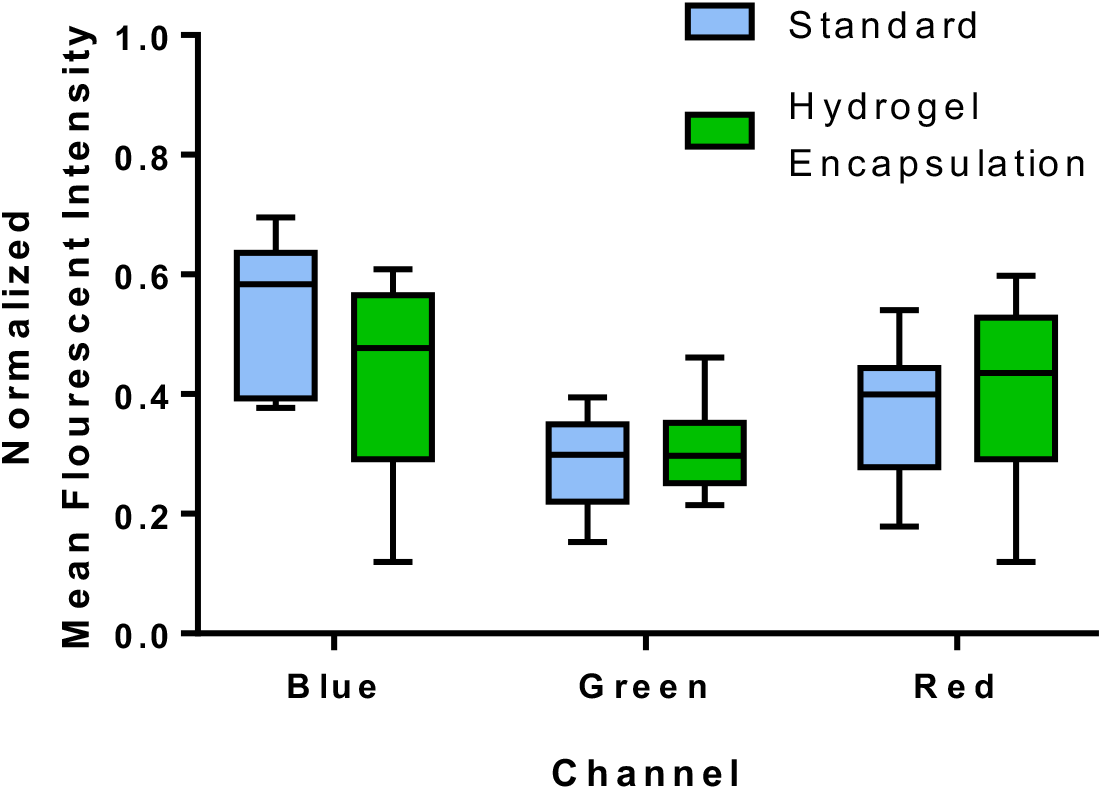
Comparison of mean fluorescent intensity. The mean fluorescent intensity (MFI) of cells following immunostaining using standard ICC and hydrogel-encapsulation. The MFI was measured from a 40×40 pixel square window surrounding each cell for each channel. Error bars represent maximum and minimum value (n=10).

### Cell Loss Comparison

We investigate cell loss during ICC resulting from convention protocol, Cytospin, and hydrogel encapsulation. Using 22RV1 prostate cancer cells as a model, we generated a 10-fold dilution series containing 10 to 10,000 cells, and then immunostained the samples using standard ICC, Cytospin, and hydrogel encapsulation. To measure cell loss, we performed triplicate experiments where cells from each specimen were enumerated by two independent reviewers before and after ICC (Figure 6). Cells stained using traditional ICC and Cytospin retained less than 50% of the cells during immunostaining regardless of the number of starting cells in the sample. In contrast, the hydrogel encapsulated cell sample retained 97-99% of input cells following immunostaining. The small deviation from ideal likely resulted from incomplete staining rather than cell loss since there are invariably a small fraction of a cell sample that will not stain. The most significant difference in cell loss was observed when fewer than 100 input cells were stained. Under these situations, almost all (>84%) cells were lost using standard ICC and Cytospin. In fact, Cytospin failed to retain any cells when there were only 10 cells in the initial sample. Together, these results show that hydrogel encapsulation permits virtually lossless immunostaining that is robust regardless of the starting number of cells in the sample.

**Fig. 6.**
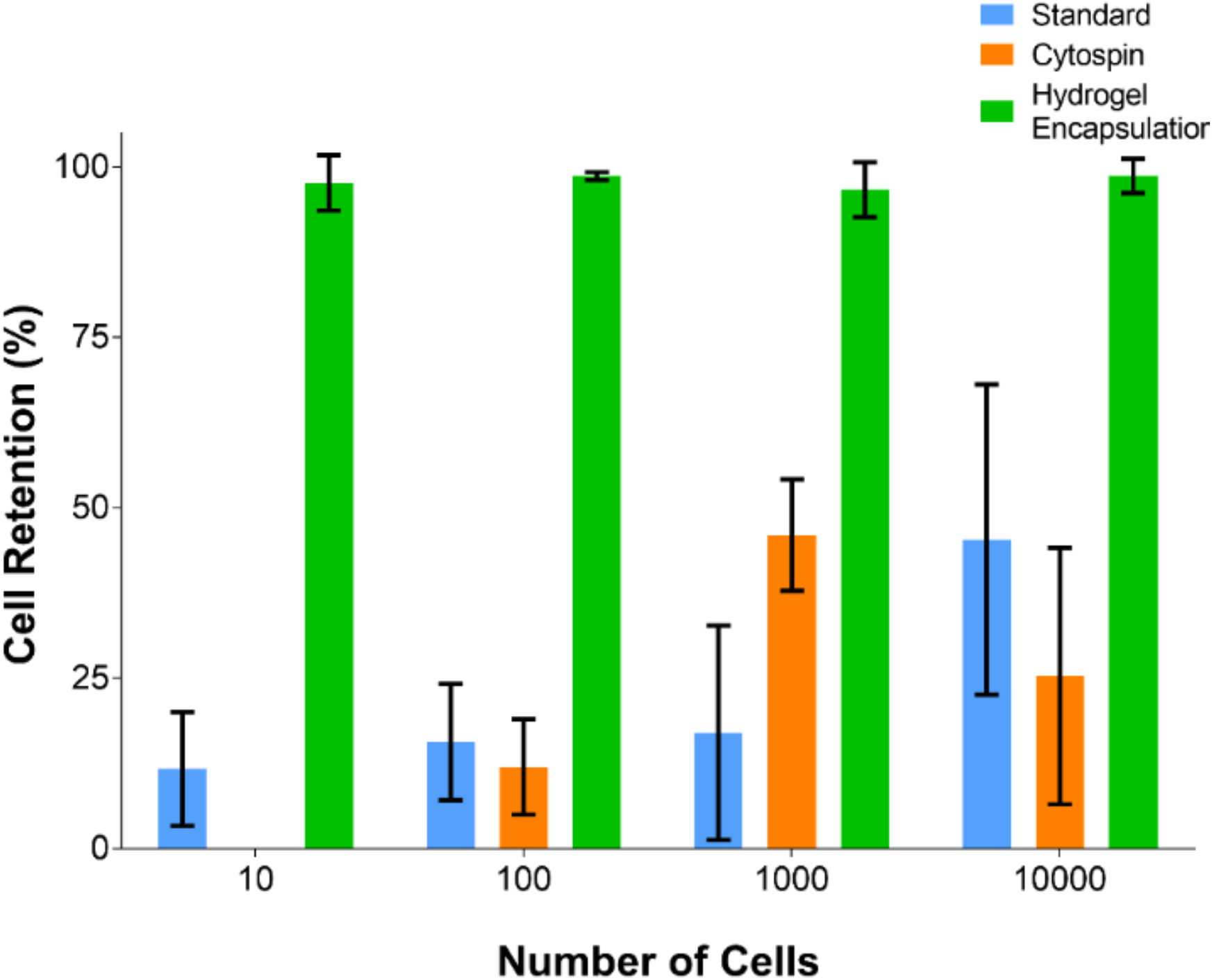
Cell retention comparison. Cell retention following immunostaining of cells using standard ICC, ICC following Cytospin, and hydrogel-encapsulation. Each experiment was performed in triplicate and cells were independently enumerated by two reviewers. Error bars rep resent the standard deviation (n=6).

## Conclusions

This technical note presents a porous hydrogel thin-film for encapsulating cells during immunocytochemistry in order to eliminate cell loss resulting from washing and centrifugation. We show that this hydrogel thin-film is permeable to immunoglobulins, stable enough to withstand pipetting, and allows immunostaining to be performed directly on standard imaging well plates. Compared to standard ICC and Cytospin methods that become highly lossy for small cell samples (<10,000 cells), this process is lossless and can be used to stain <10 cells in a well. Furthermore, this process requires only a single centrifugation step, compared to >8 steps for standard ICC, which greatly improves compatibility with robotic systems. Ultimately, this simple and novel application of hydrogels for ICC could greatly improve small cell sample biological assays, such as drug screening on primary cells and identification of rare cells, such as circulating tumor cells.

## Methods

### Chemicals and Hydrogel Preparation

The cell capture imaging reagent (LMR001) was purchased from MilliporeSigma. The paraformaldehyde (PFA), and Tween-20 were purchased from Sigma-Aldrich. Each solution was freshly prepared prior to experiments.

### Cell Culture

The cell line 22RV1 (human prostate carcinoma, ATCC CRL2505) was used for immunostaining experiments. This cell line was cultured in RPMI-1640 culture media containing 10% Fetal Bovine Serum (Gibco) and 1% penicillin-streptomycin (Gibco) at 5% CO2 at 37°C. Cells were re-suspended using 0.25% Trypsin-EDTA (Gibco), to generate a 10-fold dilution series from 10^4^−10^5^ cells per 40 μl culture media.

### Cell Encapsulation

To encapsulate the cells in hydrogel, each cell suspension and 40 μL of PBS buffer was first loaded into one well in a 384-high contrast imaging well-plate (Corning). Next, 6.5 μL of the cell capture imaging reagent solution is pipetted gently with minimal mixing. The imaging well-plate was then centrifuged for 3 minutes at 3800 rpm (Accuspin 1R, Fisher scientific), and immediately proceeded to next step.

### Photo-polymerization

To form a hydrogel thin-film, the previously prepared plate was exposed to UV light using a 375 nm UV LED (M375L3, Thorlabs) powered by a LED driver (LEDD1B, Thorlabs), or a cold cathod fluorescent lamp (CCFL) UV lamp (λ=302 nm) in a gel imaging system (Gel Doc XR+, Bio-Rad). For UV LED system, the center of the LED was aligned with the center of the each well with 0.5 mm gap and exposed for 5 seconds under drive current of 700 mA, which provides 470 mW output power. For the gel imaging system, the location of CCFL UV lamp was pre-marked, and 3 rows of well-plate were aligned on the center of each UV lamp. The exposure was controlled by Image Lab software (Bio-Rad) same as regular DNA gel imaging with a 20 second exposure.

### Cytospin Preparation

Cytospin was performed by depositing a 40 μL cell suspension directly onto a BSA-coated glass slide using a cytocentrifuge (Cytospin 2, Shadon) at 700 rpm for 3 minutes with low acceleration.

### Immunocytochemistry

Immunocytochemistry was performed by first fixing cells in 4% paraformaldehyde for 10 minutes, washing twice with PBS and permeabilizing the cells with 0.025% Tween-20 for 15 minutes. After washing the cells twice more with PBS, the cells were blocked with 3% bovine serum albumin BSA (30 min) and stained for one hour with DAPI (1 μM) for DNA, EpCAM-Alexa Fluor 488 (1:100 dilution) and Pan-Keratin-Alexa Fluor 647 (1:100 dilution). ICC was performed in parallel on matching samples that were either cytospun onto a glass slide, hydrogel-encapsulated in an imaging plate or non-encapsulated within an imaging plate. This protocol differed between cell specimens because washing of non-encapsulated cells involved adding 40 μl of PBS followed by centrifugation (3800 rpm, 3 min), washing of cytospin slides employed gentle PBS rinsing and the washing of hydrogel encapsulated cells involved adding PBS and mixing by pipette 10 times. Immunostained cells were directly imaged using both bright field and fluorescent microscopy, using a Nikon Ti-E inverted fluorescent microscope with 10x, 20x and 60x magnification with a high-resolution camera or a Zeiss laser scanning confocal microscope LSM 780 at 40x magnification.

### Cell Counting and Statistical Analysis

Both the initial (prior to plating) and final numbers of all 3 matching ICC samples were manually counted by two individuals from the obtained images using ImageJ software. Experiments were performed 3 times for each cell dilution. Results from the count were averaged and plotted using GraphPad Prism.

## Acknowledgments

We thank Peter Black for providing the 22RV1 cell line. Jeong Hyun Lee has been supported by the UBC Four Year Fellowship. Emily Park has been supported by the Michael Smith Foundation for Health Research Trainee Award. Kerryn Matthews has been supported by the MITACS Accelerate Postdoctoral Fellowship. This work has been supported by grants from Canadian Institutes of Health Research (312371, 322375, 362500, 381129), Natural Science and Engineering Research Council of Canada (2015-06541, 508392-17), and Prostate Cancer Canada (D2016-1306).

